# Generation and characterization of iPSCs from human embryonic dermal fibroblasts of a healthy donor from Siberian population

**DOI:** 10.1101/455535

**Authors:** Elena V. Grigor’eva, Tuyana B. Malankhanova, Aizhan Surumbayeva, Julia M. Minina, Elena A. Kizilova, Igor N. Lebedev, Anastasia A. Malakhova, Suren M. Zakian

## Abstract

Technology of reprogramming of somatic cells to a pluripotent state allows generating induced pluripotent stem cells (iPSCs) and carrying out a broad range of studies. iPSCs can be obtained from patients suffering from inherited diseases to model the diseases and to study their pathological mechanisms at the molecular level after iPSC differentiation in relevant cell types. Another approach to model and study inherited diseases is using iPSCs from healthy donors and genome editing tools. The approach allows generating a panel of isogenic lines, which gives new opportunities in drug screening and toxicological testing. Moreover, iPSCs and their derivatives can be further used for substitutive cell therapy and transplantology.

In this study, we generated iPSCs from human embryonic fibroblasts using episomal vectors. The lines obtained expressed pluripotency markers, had a stable karyotype – 46:XY, and did not contain episome integrations into genome. The cell lines gave rise to derivatives of three germ layers during spontaneous differentiation *in vitro* and *in vivo*.

## Derivation and characterization iMA IPSCs

Human embryonic dermal fibroblasts were reprogrammed to a pluripotent state using episomal vectors expressing reprogramming factors[1,2] (Figure 1a). The episomes were delivered into the cells by nucleofection (NHDF Nucleofector Kit, Lonza). The efficiency of nucleofection was about 50%, which was evaluated by EGFP expression in cells (Figure 1b). 23 primary clones of ESC-like morphology cells were obtained (Figure 1c). The cells intensively grew in the colonies and had dense intercellular contacts and large nuclear-cytoplasmic ratio. All clones expressed endogenous alkaline phosphatase (Figure 1d). Four clones (iMA-1T, iMA-1L, iMA-17L, iMA-23L) with well-expressed ESC-like morphology and high proliferative activity were selected for long-term culturing and subsequent characterization.

**Figure 1.**
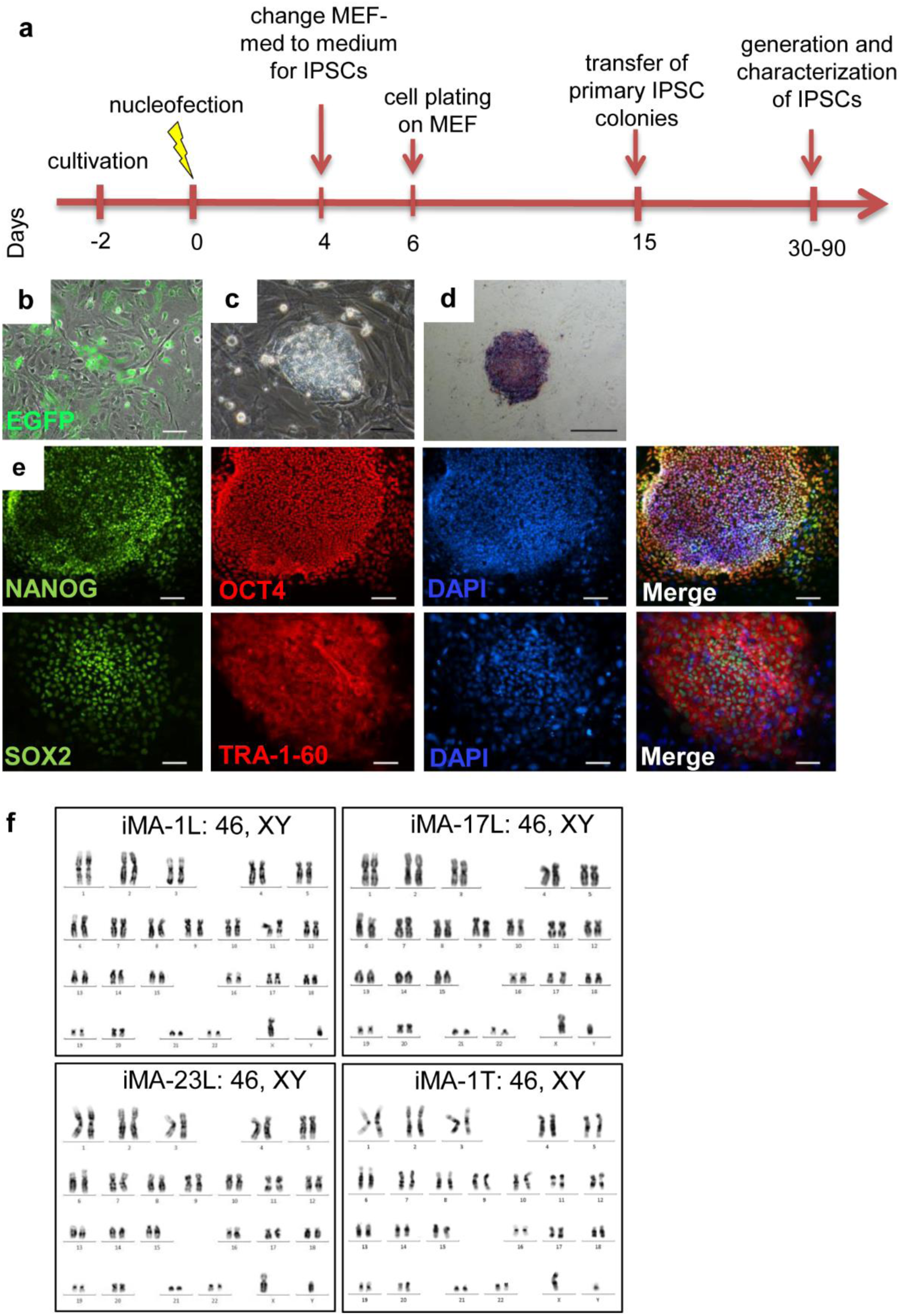
Derivation and characterization of IPSCs. (**a**) Reprogramming scheme of human fibroblasts to a pluripotent state. (**b**) GFP-positive fibroblasts on the 6 day of reprogramming. (**c**) Primary colony morphology on the 13 day of reprogramming. (**d**) Histochemical detection of endogenous alkaline phosphatase activity in iMA-1L cells, passage 1. (**e**) Immunofluorescent analysis for the pluripotency markers OCT4 (green signal), SOX2 (green signal), NANOG (red signal) and TRA-1-60 (red signal). (**f**) Karyotyping and G-band analysis show four cell IPSC lines have a normal 46, XY karyotype.

Immunofluorescent analysis for pluripotency markers in early-passage IPSCs showed the expression of OCT4, SOX2, NANOG, and TRA-1-60 in all cell lines (Figure 1e). The RT-PCR analysis also demonstrated the expression of genes, which are specific for pluripotent cells (Figure 2).

**Figure 2.**
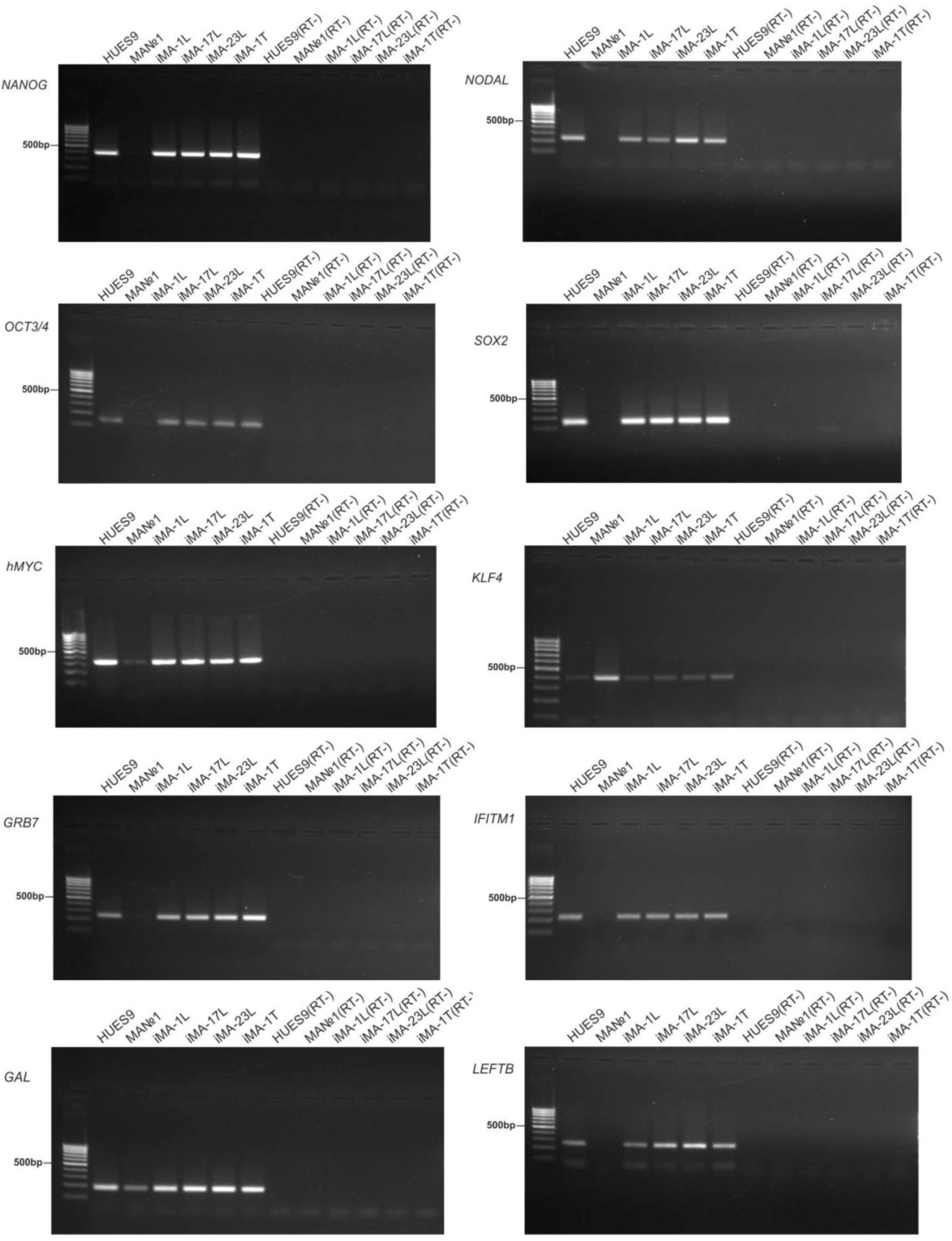
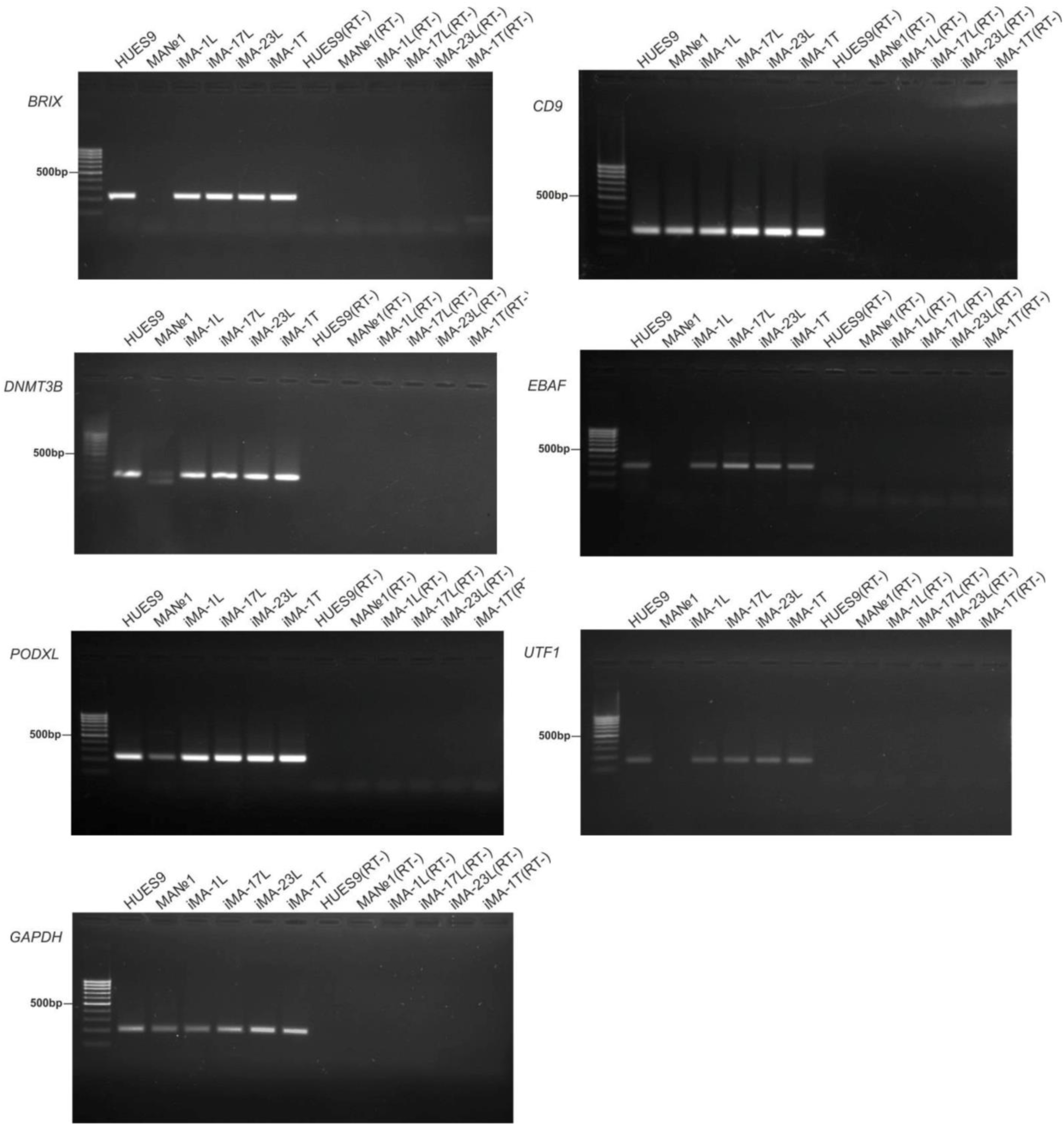
RT-PCR analysis for the expression of pluripotency markers in four IPSCs lines (iMA-1L, iMA-17L, iMA-23L, iMA-1T). MA№1 – the original embryonic dermal fibroblasts.

Karyotyping all IPSC lines showed a normal karyotype 46: XY in more than 70% of the cell population on both early (6) and late (22-24) passages (Figure 1f).

One of the general characteristics of pluripotent stem cells is the ability to produce derivatives of three germlines: ecto-, endo- and mesoderm during spontaneous differentiation. We carried out a spontaneous differentiation of IPSCs *in vitro* through embryoid bodies formation. RT-PCR analysis (Figure 3a) and immunofluorescent staining (Figure 3b) showed that there are derivatives of three germlines: ectoderm (PAX6, NF200, SOX1, GFAP), mesoderm (MSX1, αSMA, COLLAGEN I, COLLAGEN IV, FIBRONECTIN, GATA6) and endoderm (SOX17, FOXA2, GATA6).

**Figure 3.**
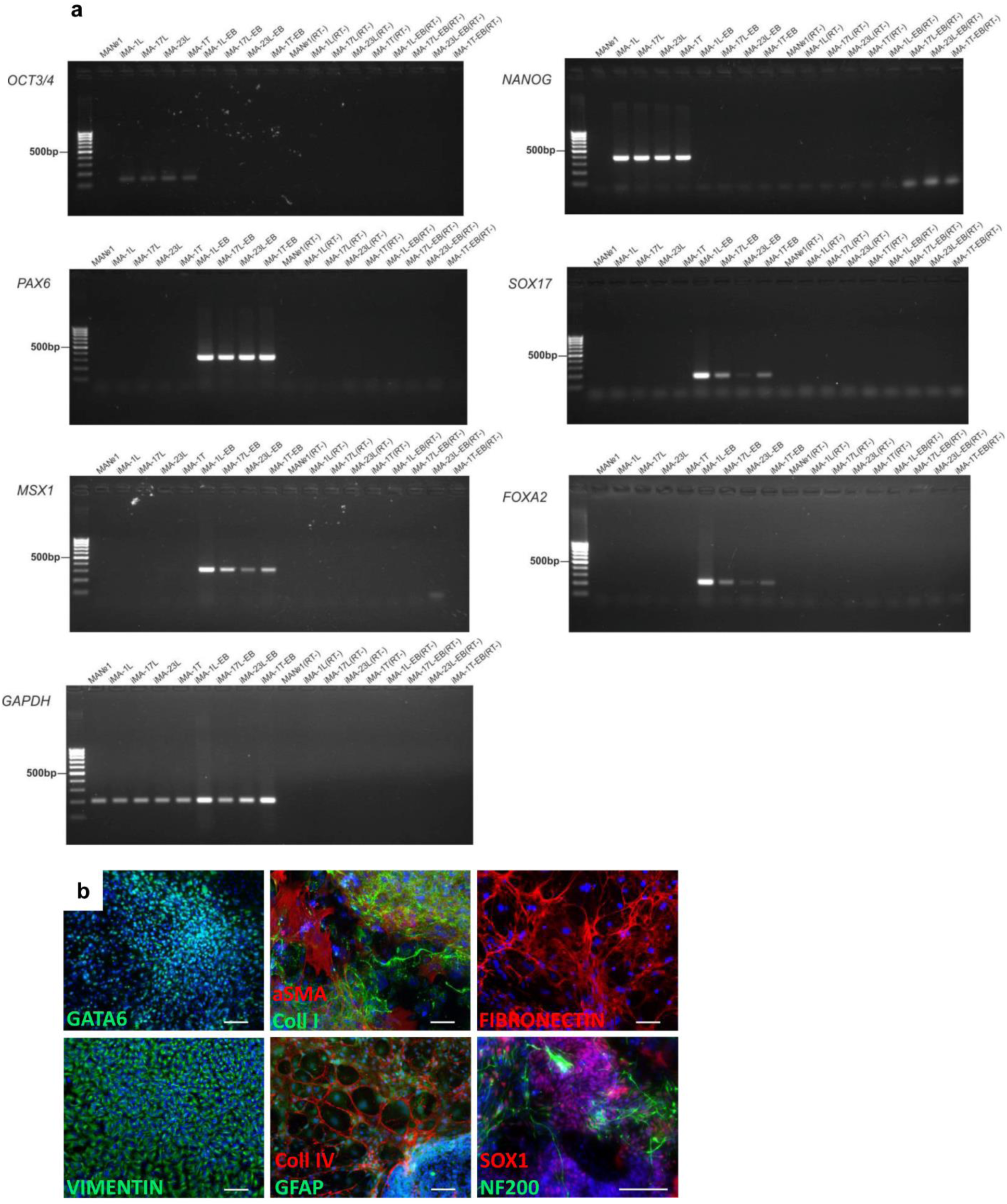
Spontaneous differentiation of IPSCs *in vitro*. (**a**) RT-PCR analysis for the expression of three germline markers. (**b**) Immunofluorescent analysis for markers of endoderm: GATA6 (green signal); mesoderm: αSMA (red signal), COLLAGEN I (green signal), COLLAGEN IV (red signal), FIBRONECTIN (red signal), VIMENTIN (red signal); ectoderm: GFAP (green signal), NF200 (green signal), SOX1 (red signal). Nuclei were stained with DAPI (blue signal). Scale bar – 100 μm, bottom right picture – 500 μm.

Spontaneous differentiation *in vivo* (teratoma assay) of 4 IPSCs lines also showed that teratomas were made up of the derivatives of three germ layers (Figure 4, Table 2). Thus, the experiments confirmed the pluripotent status of the obtained IPSCs lines.

**Figure 4.**
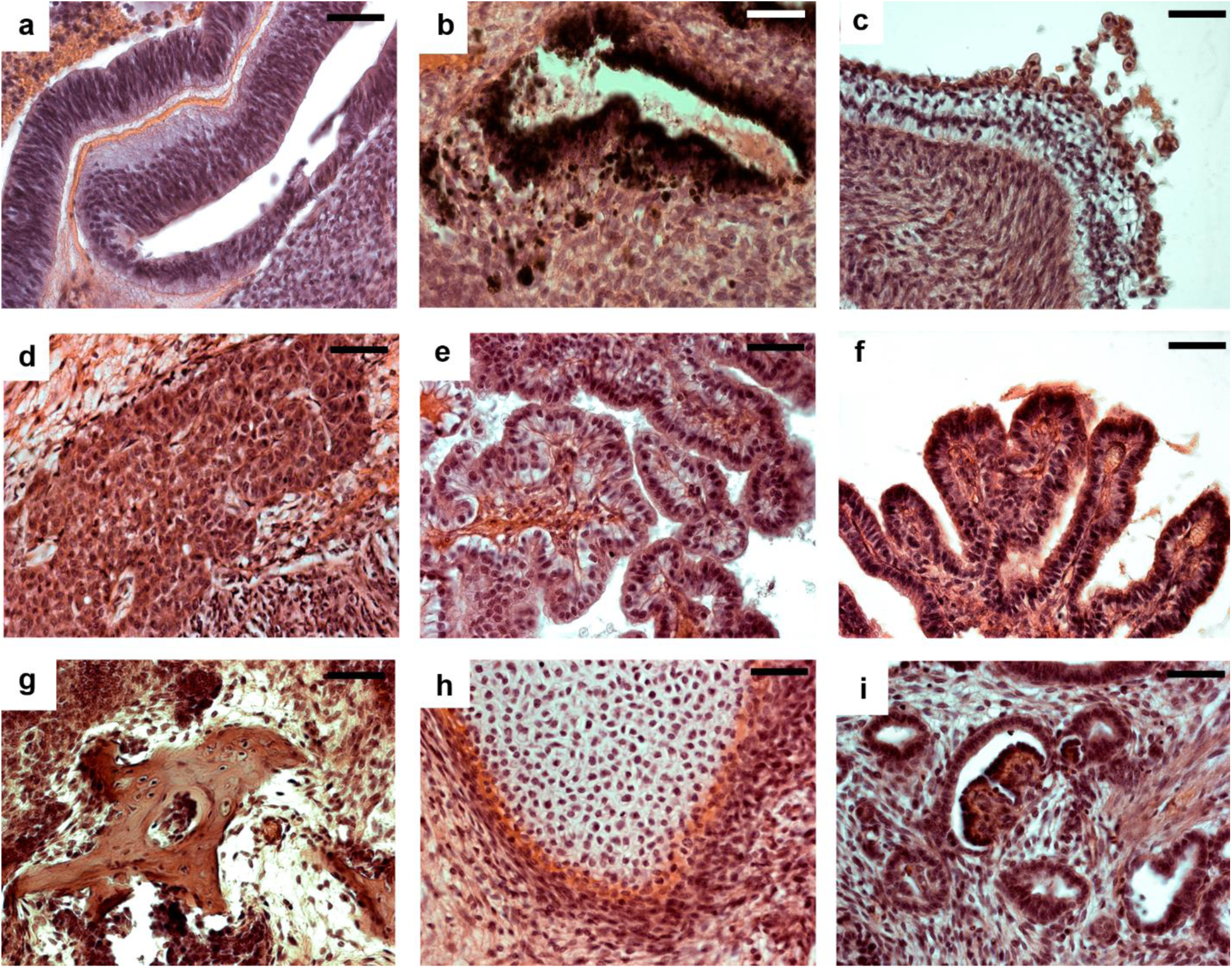
Spontaneous differentiation of IPSCs *in vivo* by teratoma assay. Derivates of the ectoderm (**a-c**), endoderm (**d-f**) and mesoderm (**g-i**). (**a**) Embryonic neuroepithelium (fold-like structures). (**b**) Pigmenteded neuroepithelium and melanocytes. (**c**) Epidermal mucoid epithelium. (**d**) Hepatocytes. (**e**, **f**) Gut epithelia. (**g**) Bone. (**h**) Cartilage. (**i**) Renal glomerullus and renal epithelia. Bar: **a**, **c**, **f**, **g** – 50 µm; **b**, **d**, **e**, **h**, **i** – 25 µm.

## Methods

### Ethical statements

The study was approved by the Scientific Ethics Committee of Research Institute of Medical Genetics, Tomsk NRMC (protocol number 106; 27th June 2017). Written informed consent was obtained from the couple. The animal tests were conducted in an SPF vivarium according to the Guidelines for Manipulations with Experimental Animals and approved by the Ethics Committee of The Federal Research Center Institute of Cytology and Genetics of the Siberian Branch of the Russian Academy of Sciences, Novosibirsk (permit No. 22.4 by 30.05.2014).

### Reprogramming of embryonic fibroblasts

The human embryonic dermal fibroblasts were reprogrammed by a set of episomal vectors (Addgene IDs #41855-58, #41813-14). Episomes were delivered into the cells using nucleofection method (NHDF Nucleofector Kit, Lonza) on the Nucleofector 2b Device (Lonza) according to the manufacturer's recommendations. The cells were maintained in MEF medium containing 50% Dulbecco’s modified Eagle’smedium (DMEM; Invitrogen), 50% F12 (Invitrogen), 10% FBS, 1xGlutaMax (Invitrogen), penicillin-streptomycin (Invitrogen). iPS cells were isolated and cultivated in a medium containing 85% Knockout DMEM (Invitrogen), 15% Knockout serum replacement (Invitrogen), 1xGlutaMax (Invitrogen), 1x nonessential amino acids (Invitrogen), and 10ng/mL basic FGF (Sigma) onto feeder cells. Feeder cells were derived from 12-day mouse embryos and treated with mitomycin C (10mg/mL; Sigma) to inhibit mitotic activity.

### Histochemical detection of endogenous alkaline phosphatase activity, immunofluorescent analysis

The detection of endogenous alkaline phosphatase and immunofluorescent staining were performed according to the previously described procedure [3]. The antibodies used for immunofluorescent staining are listed in Table 1.

**Table 1.** List of used antibodies

| Antibodies | Company | Cat. Ref. | Raised/Type | Dilution |
| --- | --- | --- | --- | --- |
| **Primary antibodies** |  |  |  |  |
| Anti SOX2 | Cell Signaling | 3579 | IgG rabbit polyclonal | 1:500 |
| Anti TRA-1-60 | Abcam | ab16288 | IgM mouse monoclonal | 1:200 |
| Anti NANOG | Abcam | ab62734 | IgG mouse monoclonal | 1:200 |
| Anti OCT4 | BD Transduction Lab | 611202 | IgG mouse monoclonal | 1:50 |
| Anti Collagen I | Abcam | ab34710 | IgG rabbit polyclonal | 1:200 |
| Anti GATA6 | Santa Cruz Biotechnology | sc-9055 | IgG rabbit polyclonal | 1:200 |
| Anti Collagen IV | LifeSpan Biosciences | LS-C79603 | IgG1 mouse monoclonal | 1:100 |
| Anti Fibronectin | Abcam | ab23750 | IgG rabbit polyclonal | 1:500 |
| Anti αSMA | DAKO | M0851 | IgG mouse monoclonal | 1:100 |
| Anti VIMENTIN | Abcam | Ab137321 | IgG rabbit polyclonal | 1:200 |
| Anti SOX1 | R&D systems | AF3369 | IgG goat polyclonal | 1:200 |
| Anti GFAP | Millipore | AB5804 | IgG rabbit polyclonal | 1:200 |
| Anti NF200 | Sigma | N4142 | IgG rabbit polyclonal | 1:1000 |
| **Secondary antibodies** |  |  |  |  |
| Alexa Fluor 488 goat anti rabbit IgG (H+L) | Thermo Fisher Scientific | A11008 | Made in goat | 1:400 |
| Alexa Fluor 568 goat anti rabbit IgG (H+L) | Thermo Fisher Scientific | A11011 | Made in goat | 1:400 |
| Alexa Fluor 488 goat anti mouse IgG (H+L) | Thermo Fisher Scientific | A11029 | Made in goat | 1:400 |
| Alexa Fluor 568 goat anti mouse IgG2a | Thermo Fisher Scientific | A21134 | Made in goat | 1:400 |
| Alexa Fluor 488 rabbit anti goat IgG (H+L) | Thermo Fisher Scientific | A11078 | Made in rabbit | 1:400 |
| Alexa Fluor 568 rabbit anti goat IgG (H+L) | Thermo Fisher Scientific | A11079 | Made in rabbit | 1:400 |
| Alexa Fluor 488 donkey anti goat IgG (H+L) | Thermo Fisher Scientific | A11055 | Made in donkey | 1:400 |
| Alexa Fluor 568 donkey anti goat IgG (H+L) | Thermo Fisher Scientific | A11057 | Made in donkey | 1:400 |
| Alexa Fluor 488 donkey anti rabbit IgG (H+L) | Molecular probes | R37118 | Made in donkey | 1:400 |

**Table 2.** Diagnostic morphotypes revealed in the histological analysis of teratomas. Ectoderm is marked by red color, mesoderm – green, endoderm – blue

| Morphotypes | iMA-1T | iMA-23L | iMA-17L | iMA-1L |
| --- | --- | --- | --- | --- |
| primitive neuroepithelium | + | + | + | + |
| premature neural (glial) tissue | + | + | + | + |
| keratinized epithelium | + |  | + |  |
| epidermal mucoid epithelia | + |  | + | + |
| melanocytes and pigmented epithelia | + | + | + | + |
| connective tissue | + | + | + | + |
| cartilage | + | + | + | + |
| bone | + | + | + | + |
| muscle | + | + | + | + |
| adipose tissue | + | + | + | + |
| erythropoietic tissue | + | + | + |  |
| lymphoid tissue |  |  |  |  |
| renal epithelia | + | + | + | + |
| ciliated epithelium |  |  |  |  |
| gut-like epithelium | + | + | + | + |
| glandular epithelium | + | + | + | + |
| hepatocytes or pancreocytes | + | + | + | + |
| number of basic diagnostic types | 15 | 13 | 15 | 13 |
| representativity, % | 88% | 76% | 88% | 76% |

### Cytogenetic analysis

IPSCs were placed onto 60 mm culture Petri dishes coated MEF feeder layer 2-3 days prior to fixation. On the fixation day, the medium was refreshed and 0.05 μg/ml of KaryoMAX Colcemid (Life Technologies) was added for 3 h. The cells were then washed with PBS and TrypLE solution was added for 3 min. Without removing TrypLE, 0.28% KCl (3 ml) was added to the cells in Petri dish and incubated for 18 min at 37 °C. Then, 2-3 drops of Carnoy fixation (3:1 methanol:acetic acid) were added for prefixation. Cells were carefully suspended, transferred into a tube and centrifuged for 5 min at 1400 rpm. The supernatant was removed; 1 ml of ice-cold Carnoy fixator was gently poured onto the precipitate and incubated for 20 min on ice. The cells were carefully pipetted and centrifuged for 5 min at 1400 rpm. The supernatant was discarded; the cells were resuspended into 1 ml of Carnoy fixation and incubated for 10 min on ice. Centrifugation was repeated for 5 min at 1400 rpm, fresh ice-cold Carnoy was added to the cells. 50-70 μl of the cell suspension was dripped onto the cooled wet slides and dried under the fan warm air. For karyotype analysis the slides with metaphase spreads were stained by DAPI (4′,6-diamidino-2-phenylindole) (50 ng/ml). 100 metaphases were analyzed.

### Spontaneous differentiation of IPSCs in vitro

IPSCs were desegregated with 0.15% collagenase and plated onto Petri dishes coated with 1% agarose in the IPSCs medium without bFGF. On the next day, cells were centrifuged at 500 rpm and resuspended in DMEM/F12 (1:1) with 10% FBS (Life Technologies), 1x GlutaMAX (Life Technologies), 1x pen/strep (Lonza). The embryoid bodies were cultured for 14 days in suspension and then were plated onto Petri dishes coated with 0.1% gelatin, where they were spread out and cultured for another 7-14 days. The differentiated cells were analyzed by immunofluorescent analysis or RT-PCR.

### RNA isolation, RT-PCR analysis of IPSCs and their differentiated derivatives

The isolation of RNA and RT-PCR were performed according to the previously described procedures[3,4]. The primer sequences used in RT-PCR are given in Supplement Table 3.

**Table 3.**
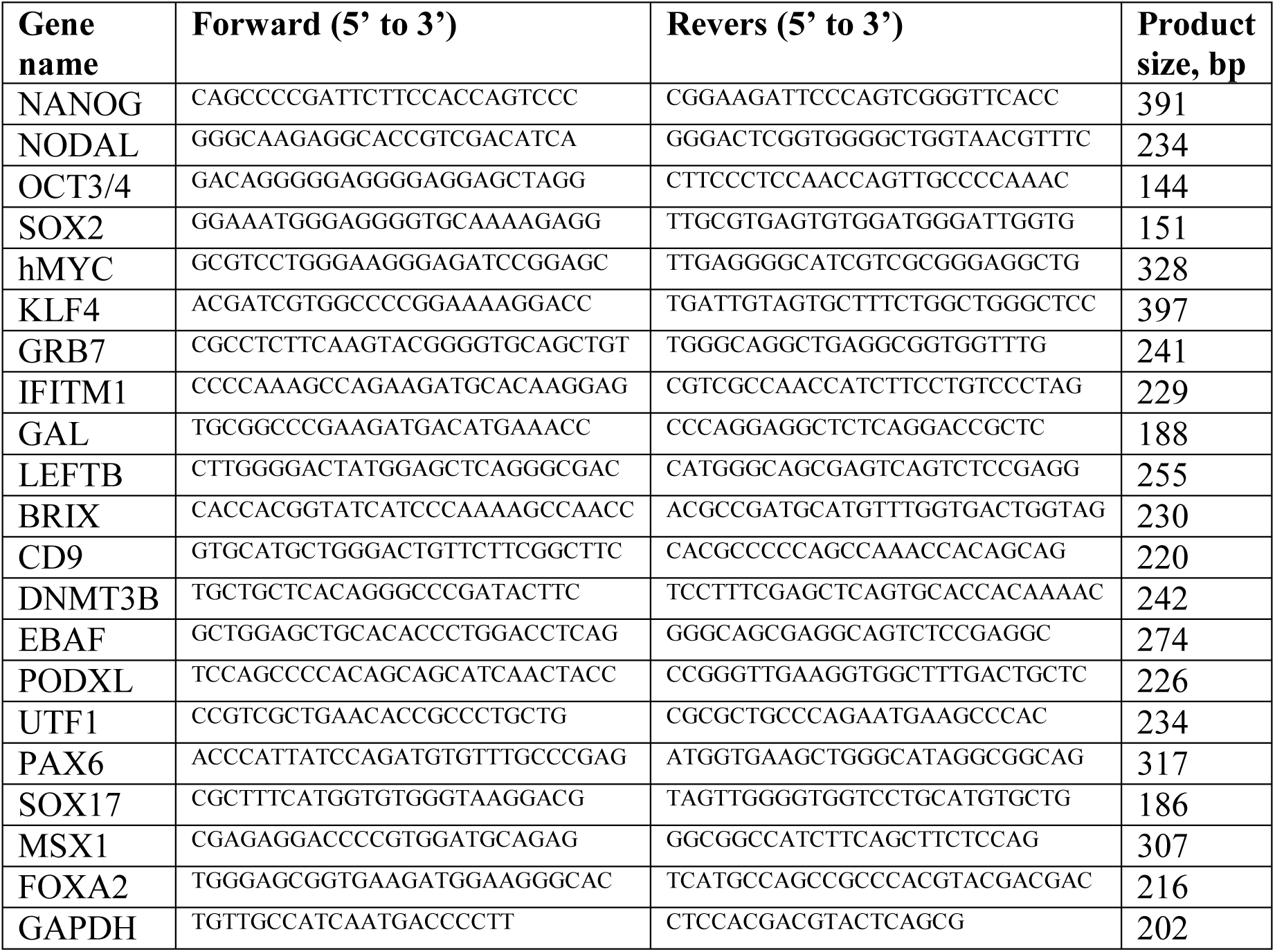
List of used primer sequences

### Teratoma assay

The teratoma assay was carried out on the basis of the SPF-vivarium of the Federal Research Center Institute of Cytology and Genetics of the Siberian Branch of the Russian Academy of Sciences (http://spf.bionet.nsc.ru).

Spontaneous differentiation *in vivo* was performed by IPSCs suspension injection (iMA-1T, iMA-1L, iMA-17L, iMA-23L) in SCID mice; 4 animals per IPSCs line. Cell suspension (3-5×10^6^ cells in 70 μl Matrigel-GFR matrix) were injected in the scruff of the neck behind the ears. After 6-8 weeks, tumor material was removed from the animals and histological analysis was performed. The preparation and histological analysis of the tumor material was carried out according to routine protocols, as described earlier[5]. Images were captured and analyzed using Axioscop2+ supplied with CCD camera (AxioCam HRc) and software, AxioVision, in the Interinstitutional Shared Center for Microscopic Analysis of Biological Objects SB RAS (www.bionet.nsc.ru/labs/viv/index.php?id=113).

